# Reprogrammed Human Lateral Ganglionic Eminence Precursors Generate Striatal Neurons and Restore Motor Function in a Rat Model of Huntington’s Disease

**DOI:** 10.1101/2024.10.09.617504

**Authors:** Amy McCaughey-Chapman, Anne Lieke Burgers, Catharina Combrinck, Laura Marriott, David Gordon, Bronwen Connor

**Affiliations:** Department of Pharmacology and Clinical Pharmacology, Centre for Brain Research, School of Medical Science, Faculty of Medical and Health Sciences, University of Auckland, Auckland, New Zealand

**Keywords:** Huntington’s disease, lateral ganglionic eminence precursor cells, striatum, direct reprogramming, cell replacement therapy

## Abstract

**Background:** Huntington’s disease (HD) is a genetic neurological disorder predominantly characterised by the progressive loss of GABAergic medium spiny neurons in the striatum resulting in motor dysfunction. One potential strategy for the treatment of HD is the development of cell replacement therapies to restore neuronal circuitry and function by the replacement of lost neurons. We propose the generation of lineage-specific human lateral ganglionic eminence precursors (hiLGEP) using direct reprogramming technology provides a novel and clinically viable cell source for cell replacement therapy for HD.

**Methods:** hiLGEPs were derived by direct reprogramming of adult human dermal fibroblasts (aHDFs) using chemically modified mRNA (cmRNA) and a defined reprogramming medium. hiLGEPs were differentiated *in vitro* using an optimised striatal differentiation medium. Acquisition of a striatal precursor and neural cell fate was assessed through gene expression and immunocytochemical analysis of key markers. hiLGEP-derived striatal neuron functionality *in vitro* was demonstrated by calcium imaging using Cal-520. To investigate the ability for hiLGEP to survive, differentiate and functionally integrate *in vivo*, we transplanted hiLGEPs into the striatum of quinolinic acid (QA)-lesioned rats and performed behavioural assessment using the cylinder test over the course of 14 weeks. Survival and differentiation of hiLGEPs was assessed at 8 and 14-weeks post-transplant by immunohistochemical analysis.

**Results:** We demonstrate the capability to generate hiLGEPs from aHDFs using cmRNA encoding the pro-neural genes *SOX2* and *PAX6*, combined with a reprogramming medium containing Gö6983, Y-27632, N-2 and Activin A. hiLGEPs generated functional DARPP32+ neurons following 14 days of culture in BrainPhys™ media supplemented with dorsomorphin and Activin A. We investigated the ability for hiLGEPs to survive transplantation, differentiate to medium spiny-like striatal neurons and improve motor function in the QA lesion rat model of HD. Fourteen weeks after transplantation, we observed STEM121+ neurons co-expressing MAP2, DARPP32, GAD65/67, or GABA. Rats transplanted with hiLGEPs also demonstrated reduction in motor function impairment as determined by spontaneous exploratory forelimb use when compared to saline transplanted animals.

**Conclusion:** This study provides proof-of-concept and demonstrates for the first time that aHDFs can be directly reprogrammed to hiLGEPs which survive transplantation, undergo neuronal differentiation to generate medium spiny-like striatal neurons, and reduce functional impairment in the QA lesion rat model of HD.

**Significance statement:** The present study reports for the first time that human lateral ganglionic eminence precursor (hiLGEP) cells directly reprogrammed from adult human fibroblasts using chemically modified mRNA can survive transplantation into the quinolinic acid-lesioned rat striatum and generate medium spiny striatal neurons. Most importantly, the authors show that transplantation of directly reprogrammed hiLGEPs restores motor function impairment by 14 weeks post-transplantation. This work provides proof of concept and demonstrates that directly reprogrammed hiLGEPs offer an effective and clinically viable cell source for cell replacement therapy to treat Huntington’s disease.

## Introduction

Huntington’s disease (HD) is a genetic neurological disorder caused by an expansion mutation of the trinucleotide (CAG) repeat in exon 1 of the *HTT* (IT15) gene, encoding a 350-kDa protein termed Huntingtin (HTT). The disease is inherited in an autosomal dominant manner and shows a prevalence of about 1 to 15,000 individuals. HD is characterised by neuronal cell loss mainly in the caudate nucleus, putamen, and the cerebral cortex. In later stages, areas such as the hippocampus and hypothalamus are affected [1]. Predominant degeneration of medium-sized spiny projection neurons (MSNs) results in motor dysfunction together with cognitive and psychiatric disturbances.

Current treatment options for HD are severely limited. While some of the behavioural symptoms of HD respond to psychiatric treatments and several drugs are available to reduce the impact of chorea, other motor symptoms and the cognitive symptoms of HD are currently not treatable [2]. Cell transplantation is a viable option for the treatment of HD with the aim of reconstructing the damaged neural circuitry by replacing cells lost to the disease process, with the expectation that the donor cells will reconnect to remaining host neural networks to repair connectivity. Through genetic testing early therapeutic intervention is possible allowing for transplantation of replacement MSNs prior to extensive degeneration with the aim of maintaining the corticostriatal circuitry. Both rodent and primate HD studies have demonstrated that reconstruction of the corticostriatal circuitry following cell transplantation can alleviate both motor and cognitive deficits observed in HD [3–5]. Human studies have demonstrated safety and feasibility, with proof-of-concept evidence to suggest that human foetal grafts can improve function, although solid evidence of efficacy is still required [6–8]. However, for cell transplantation therapy to be a viable therapeutic option for HD patients, one of the main issues that needs to be addressed is the identification of an ethically and technically viable source of donor cells other than human foetal tissue.

In searching for an alternative donor cell source attention has fallen on the potential use of human-derived stem cells including human embryonic stem cells (hESCs) or human induced pluripotent stem cells (iPSC) [9]. Previous studies have demonstrated that hESC-derived neural stem cells (NSC) transplanted into the quinolinic acid (QA) lesion model of HD survive and generate region-specific neurons expressing markers of MSNs [10–14]. To increase lineage specificity and enhance differentiation into MSNs, several groups differentiated hESCs into lateral ganglionic eminence precursors (LGEPs) [15–19]. These studies reported increased generation of MSNs following transplantation of LGEPs into the QA lesioned striatum demonstrating the requirement to drive hESCs to a lineage-specific precursor fate before transplantation. Several studies have also demonstrated the potential use of iPSC-derived NSCs as a cellular source for transplantation therapy for HD [20, 21], with An and colleagues [21] demonstrating the capability to transplant genetically corrected HD patient-derived cells.

An exciting alternative cell source for the treatment of HD is the generation of human induced neural precursor cells (hiNPCs) through direct cell reprogramming [22]. We have developed a direct reprogramming strategy utilising the neural promoting transcription factors *SOX2* and *PAX6* to reprogram adult human dermal fibroblasts (aHDFs) to a neural precursor-like state [22, 23]. Transient over-expression of *SOX2* and *PAX6* is achieved by the use of chemically modified mRNA (cmRNA) resulting in the generation of hiNPCs which can be differentiated to a GABAergic or glutamatergic neuronal lineage in conjunction with astrocytes [22]. The use of cmRNA for direct cell reprogramming is ideal for cell transplantation as it is an efficient gene delivery system with a high safety profile required for the generation of neural cells for potential clinical use [24, 25].

To transfer our direct cell reprogramming technology for use in cell transplantation therapy, the current study demonstrates the ability to generate human induced lateral ganglionic eminence precursor (hiLGEP) cells using cmRNA direct reprogramming. This study provides proof of concept and demonstrates for the first time that aHDFs can be directly reprogrammed to hiLGEPs which survive transplantation, undergo neuronal differentiation to generate a population of cells expressing markers characteristic of striatal neurons, and reduce functional impairment in the QA lesion rat model of HD.

## Materials and Methods

### Adult human dermal fibroblast direct-to-induced lateral ganglionic eminence precursor cell reprogramming and differentiation

Human induced lateral ganglionic eminence precursor (hiLGEP) cells were generated from adult human dermal fibroblast (aHDF) cell lines (1507: Male Caucasian, 50 years old, facial tissue; 1838: Male Caucasian, 50 years old, facial tissue; 2116: Female Caucasian, 35 years old, abdominal tissue; 2298: Female Caucasian, 33 years old, abdominal tissue; Cell Applications Inc; 31037: Male Caucasian, 30 years old; 30625: Male Caucasian, 76 years old; NINDS). aHDF cells were cultured in Dulbecco’s modified eagle medium (DMEM; Thermo Fisher Scientific) containing 10% fetal bovine serum (FBS; Thermo Fisher Scientific). aHDFs were induced to a lateral ganglionic eminence precursor (LGEP) fate by transient over-expression of the pro-neural genes *SOX2* and *PAX6* using chemically modified mRNA (cmRNA; Ethris GmbH, Munich, Germany) [22]. The aHDFs were seeded at 300,000 cells/well in Nunc-coated 6 well plates (Nunc) and transfected with 2.5µg of each *SOX2* and *PAX6* cmRNA using Lipofectamine RNAiMAX (Invitrogen) transfection reagent. Five-hour transfections were conducted over four consecutive days. The cells were reprogrammed under normoxia in either a Neurobasal-A (NBA; Gibco) or a BrainPhys™-based (Stem Cell Technologies) reprogramming medium containing 1mM valproic acid (Sigma Aldrich), 1% penicillin-streptomycin-glutamine (Gibco), 2% B-27 without retinoic acid (Gibco), 20ng/ml EGF (Prospec Bio), 20ng/ml FGF2 (Prospec Bio), 2µg/ml heparin (Sigma Aldrich), 1% N-2 supplement (Gibco), 5µM Gö6983 (Abcam), 10µM Y-27632 (Abcam), and 10µM retinoic acid (Sigma Aldrich). The cells were passaged at Day 7 of reprogramming using 0.05% trypsin-EDTA (400µL/well; Gibco) and trypsin inhibitor (400µL/well; Sigma) and re-seeded to 300,000 cells/well on 6 well plates. Activin A (25ng/mL; Prospec Bio) was added to the reprogramming medium from Day 7 to Day 14 of reprogramming (Figure 1). Typically, the number of cells obtained by Day 14 of reprogramming was double the number of aHDFs seeded for transfection. The gene expression of *SOX2* and *PAX6* was seen to reduce gradually over the course of reprogramming (Supplementary Figure 1A), with no SOX2 protein expression and little PAX6 protein expression found in hiLGEPs (Supplementary Figure 1B). No SOX2 and PAX6 protein expression was seen in hiLGEP-derived cells at Day 14 of differentiation (Supplementary Figure 1C).

**Figure 1:**
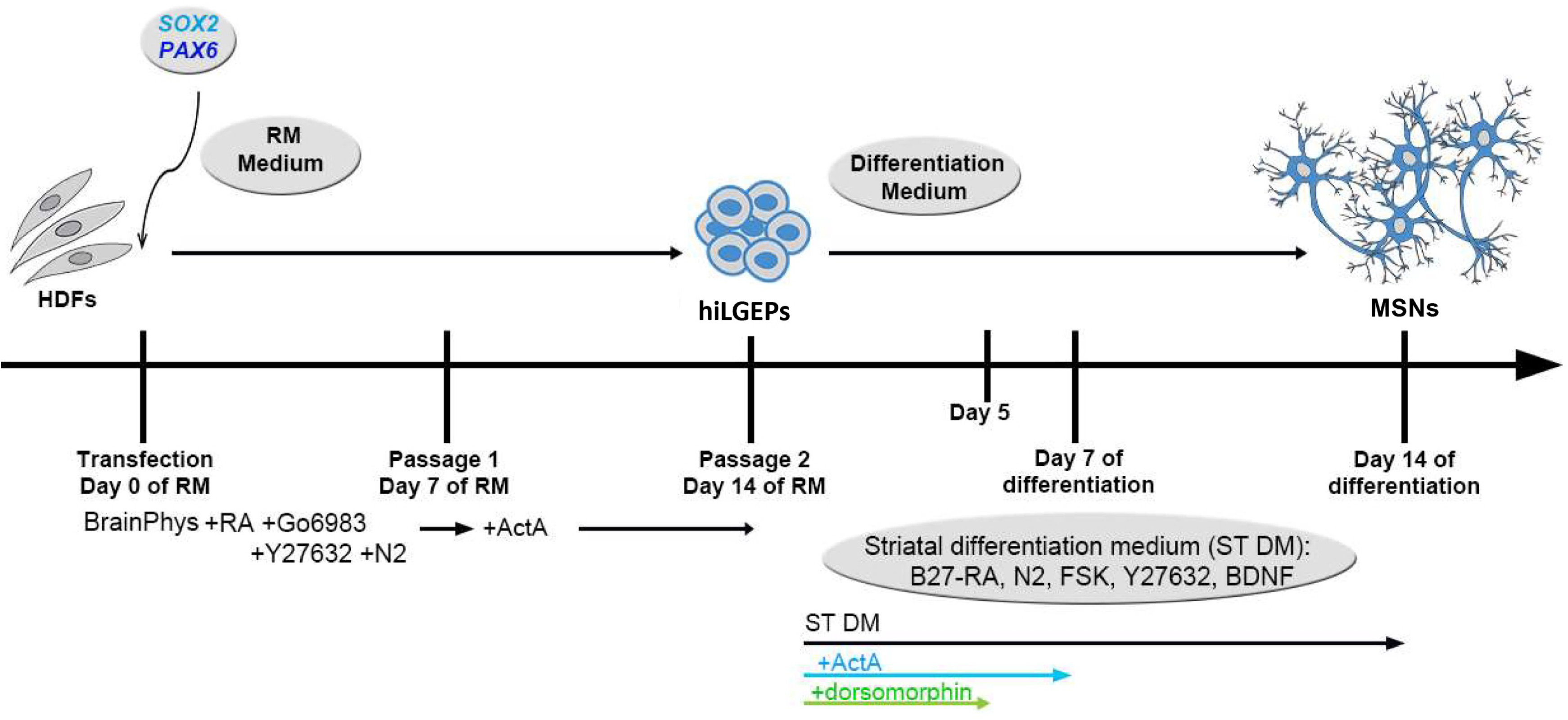
Schematic diagram of *SOX2/PAX6* chemically modified mRNA direct reprogramming of adult human dermal fibroblasts (HDFs) to generate human induced lateral ganglionic eminence precursors (hiLGEPs). hiLGEPs are then differentiated in striatal differentiation media for 14 days to generate medium spiny neurons (MSNs). RM = reprogramming; RA = retinoic acid; ActA = Activin A; FSK = forskolin.

Prior to reprogramming, a set of aHDFs were transduced with a lentiviral vector expressing zsGreen (LV-zsGreen) to enable visualisation of the cells following transplantation. Three days after transduction, approximately 80% of aHDFs expressed zsGreen protein. The aHDFs were then transfected with *SOX2* and *PAX6* mRNA and reprogrammed for 14 days at which time they were collected for transplantation into the QA lesioned rat striatum. The zsGreen-labelled cells were used for animals culled at 8 weeks post-transplantation only, unlabelled hiLGEPs were used for animals culled at 14 weeks post-transplantation.

At Day 14 of reprogramming, hiLGEPs were collected either for transplantation using StemPro™ Accutase™ (Gibco) or were processed for RT-qPCR and immunocytochemistry to confirm the acquisition of a LGEP cell fate. An additional subset of cells were plated out for differentiation at 60,000 cells/well on GelTrex-coated glass coverslips and cultured in either NBA-based or BrainPhys™-based striatal differentiation media containing 1% penicillin-streptomycin-glutamine, 2% B-27 supplement without retinoic acid, 1% N-2 supplement, 10µM Y-27632, 10µM forskolin (Abcam), 30ng/mL BDNF (Prospec Bio) and for the first 5 days, 1µM dorsomorphin (Abcam) with or without 5µM Gö6983, and the first 7 days 25ng/mL Activin A. After 14 days of differentiation, the cells were fixed with 4% paraformaldehyde at 4⁰C and processed for immunocytochemistry.

### Quantitative RT-PCR

Total RNA was isolated from hiLGEPs at 14 days of reprogramming and the originating aHDF cell line using the Nucleospin RNA kit (Macherey Nagel). cDNA was synthesised from total RNA using Superscript IV reverse transcriptase (Thermo Fisher Scientific). Duplex qPCR reactions were performed using the TaqMan® system (Applied Biosystems) with ribosomal 18S rRNA as the internal standard and an equivalent of 4-10ng RNA per reaction, in triplicate. Gene expression was normalised to ribosomal 18S rRNA as the internal standard. Gene expression was presented as fold change relative to aHDFs using the ΔΔCt method.

### Immunocytochemistry

The cells were permeabilised in phosphate-buffered saline with 0.5% Triton X-100 for 5 minutes and then incubated in primary antibody-containing immunobuffer overnight at 4°C. The following human-specific primary antibodies were used: CTIP2 (1:250, Abcam), FOXP1 (1:100, R&D), FOXP2 (1:500, Abcam), MEIS2 (1:500, Abcam), TUJ1 (1:500, Abcam), DARPP32 (1:500, Invitrogen), GABA (1:500, Invitrogen), GAD65/67 (1:500, Abcam), SOX2 (1:250, Abcam), PAX6 (1:500, BioLegend), MAP2 (1:1,000, Abcam), SYN1 (1:250, Abcam) and PSD-95 (1:250, Synaptic Systems). The species-appropriate Alexa Fluor™ secondary antibodies (1:500; Invitrogen) were used for visualisation of the primary antibody by incubation for 1 hour at room temperature. DAPI included in Prolong Diamond antifade mountant (Invitrogen) was used to confirm individual cell nuclei. Images were captured using an inverted Nikon TE2000E fluorescence microscope equipped with a DS-Ri2 camera. Quantification of the number of CTIP2+, FOXP1+, FOXP2+, MEIS2+ hiLGEPs and TUJ1+, DARPP32+, GABA+ or GAD+ hiLGEP-derived neurons was undertaken manually in ImageJ as a proportion of DAPI+ cells from a minimum of 500 DAPI+ cells. MAP2/SYN1 and MAP2/PSD-95 images were captured using an inverted LSM 800 scanning laser confocal microscope (Zeiss) and a 40x oil objective.

### MACS® MicroBead magnetic cell sorting

hiLGEPs were generated as described above by reprogramming for 14 days in either NBA-based reprogramming media with GYN and Activin A, BrainPhys™-based reprogramming media with Activin A and no GYN or in BrainPhys™-based reprogramming media with GYN and Activin A. hiLGEPs were then plated out for differentiation at 80,000 cells/well on GelTrex-coated 24-well, black wall, glass bottom Sensoplates (Greiner Bio-One GmbH) and cultured in either NBA-based striatal differentiation media or BrainPhys™-based striatal differentiation media for 14 days as described above (12 replicate wells per condition per line with 3 cell lines). After 14 days of differentiation, the cells were fixed in 4% PFA for 10 minutes at room temperature and the collected with Accutase. The 12 replicate wells per media condition and line were pooled into a single Falcon tube, centrifuged at 350g for 5 minutes, resuspended in 200µL of PBS with 0.5% BSA containing 2µL of PGP9.5 (Abcam, mouse IgG) and incubated at 4°C overnight. The next day the cells were washed by addition of 1mL of PBS with 0.5% BSA and centrifuged at 300g for 10 minutes. The supernatant was aspirated, and the cell pellet resuspended in 80µL of PBS with 0.5% BSA. 20µL of anti-mouse IgG MicroBeads were added to the cell suspension and incubated at 4°C for 15 minutes. The cells were then washed by addition of 1mL of PBS with 0.5% BSA and centrifuged at 300g for 10 minutes. The supernatant was again aspirated, and the cell pellet resuspended in 500µL of PBS with 0.5% BSA. The unlabelled and labelled cells were then isolated by magnetic separation using a MS column. The MS column was placed into the magnetic field of a MACS Separator and rinsed with 500µL of PBS with 0.5% BSA. The cell suspension was applied to the column and the flow-through of unlabelled cells collected. The column was rinsed three times with 500µL of PBS with 0.5% BSA and the flow-through added to the unlabelled cell effluent. The column was then removed from the separator, placed in a Falcon tube, 1mL of PBS with 0.5% BSA was applied to the column and the labelled cells were immediately flushed out by firmly pushing the plunger through the column. The number of unlabelled and labelled cells for each sample were counted using a Haemocytometer and the proportion of labelled cells calculated as a percentage of total cells. An average percentage of PGP9.5 labelled cells was determined for each media condition from n=3 independent cell lines.

### Live-cell calcium imaging

hiLGEPs were plated out for differentiation at 80,000 cells/well on GelTrex-coated 24-well, black wall, glass bottom Sensoplates (Greiner Bio-One GmbH) and cultured in BrainPhys™-based striatal differentiation media for 14 days as described above. The cells were loaded with a working solution comprising of BrainPhys™ media, 5μM of Cal-520 AM (Abcam) with 0.04% Pluronic F-127 (Invitrogen) and incubated at 37°C for 1 hour, followed by incubation at room temperature for 30 minutes. The Cal-520 AM dye working solution was replaced with Hank’s Balanced Salt Solution buffer without phenol red (Gibco) containing 1mM Probenecid and 20mM 4-(2-hydroxyethyl)-1-piperazineethanesulfonic acid (HEPES; Gibco). Cells were imaged at Ex/Em = 490/525nm for Cal-520 AM fluorescence intensity for a total of 120 seconds, with or without glutamate treatment at 12.5, 25, 37.5, and 50µM. Timelapse recordings were captured on a Nikon TE2000E inverted microscope equipped with a Nikon Digital Sight DS-Ri2 CMOS sensor colour camera using NIS Elements AR (Advanced Research) software. Live-cell calcium imaging analysis was conducted using the Time Series Analyser V3 plugin in FIJI. Average florescence intensity values for 75 regions of interest were recorded from 2 replicates. Fluorescence was measured as the percentage increase in average fluorescence intensity relative to baseline fluorescence at time 0 and then normalised to the average fluorescence intensity measured in response to no addition of glutamate at each time point.

### Animals

Adult male Sprague-Dawley rats (University of Auckland Vernon Jansen Unit) 250 to 350 g (8 weeks old) at the time of quinolinic acid (QA) lesions were used in this study. All procedures strictly complied with the University of Auckland Animal Ethics Guidelines, in accordance with the New Zealand Animal Welfare Act 1999 and international ethical guidelines. All efforts were made to minimize the number of animals used and their suffering. The rats were randomly allocated into the following treatment groups: hiLGEP transplants – 8 weeks (n = 5); hiLGEP transplants – 14 weeks (n = 15); sham transplants (0.9% sterile saline, 8 weeks: n = 5; 14 weeks: n = 15). The number of animals included in the study and assigned to each group was based off our previous animal studies and to allow for unplanned casualties. Animals culled at 8 weeks post-transplant were used for immunohistochemistry (n = 5 animals per group). Animals culled at 14 weeks post-transplant were used for both behavioural analysis and immunohistochemistry (n = 15 per group). The rats were group-housed in a temperature-and humidity-controlled room on a 12-hour light/dark cycle. Food and water were available *ad libitum* throughout the whole study. 48 hours before transplant surgery, all rats received an intraperitoneal injection of Sandimmun (20mg/kg Cyclosporine; supplied by the Vernon Jansen Unit, University of Auckland). Following the transplant surgery, Sandimmun injections continued thrice weekly for the duration of the study.

### Surgical procedures

All surgeries were performed using isoflurane anaesthetic (induction 5% isoflurane and a flow rate of 2 L/min O2; maintenance 1.5% isoflurane and a flow rate of 1.5 L/min O2). All rats received a unilateral intrastriatal infusion of QA (50 nmol, 400 nl; 100 nl/min with a 32G Hamilton syringe controlled by a WPI UltraMicroPump II) at the following stereotaxic coordinates: +0.5 mm anterior-posterior (AP), -2.7 mm medial-lateral (ML), relative to bregma and -5.0 mm dorsal-ventral (DV) from the dural surface [12, 26, 27]. Injections of hiLGEPs (∼250,000 viable cells per animal) or 0.9% sterile saline were performed 21 days after QA lesioning into two adjacent sites in the lesioned striatum (∼62,500 viable cells/µl: 2 µl per injection site; 400 nl/min with a 26G Hamilton syringe) at the following stereotaxic coordinates: +0.3 mm AP, -2.5 mm ML, at -5.0 and -4.0 mm DV [12, 26, 27]. Rats underwent surgeries in no particular order, in fact in a random order to limit confounder effects.

### Spontaneous exploratory forelimb use

The rats were placed in a Plexiglas cylinder (20 cm diameter) and their behavior videotaped for 5 minutes. Rats underwent the test in no particular order to avoid confounder effects. Baseline tests of motor function were obtained prior to QA lesioning. Motor function was also assessed 2 weeks following QA lesioning. Following transplantation, the rats were assessed at 2-, 4-, 12- and 14-weeks post-transplantation [Figure 5A]. Spontaneous exploratory forelimb use was scored by an experimenter blind to the condition of the animals during slow-motion feedback of the videotaped sessions using forelimb asymmetry analysis as described [28, 29]. Forelimb use was assessed as a single asymmetry score representing the overall percentage ipsilateral forepaw use for wall placement during exploratory rearing over a 5-minute trial period and the first 20 rears [26, 27]. Rats that failed to respond to the QA lesion by predominantly using the forelimb contralateral to the lesion were removed from all analysis. Analysis was undertaken by an investigator blind to the group allocation.

### Immunohistochemical analysis

Rats were culled 8 or 14 weeks after transplantation with sodium pentobarbital (120mg/kg i.p.) followed by transcardial perfusion with 0.9% saline and 4% paraformaldehyde. Brains were cryoprotected in 30% sucrose before sectioning coronally at 40 µm on a HM450 sliding microtome (Microm International GmbH, Walldorf, Germany) or Leica CM3050 cryostat. Eight sets of sections were collected from each brain (distance 320 µm between consecutive sections in each set) and stored at -20⁰C.

Fluorescent immunohistochemistry was performed on free-floating coronal sections from each animal using antibodies against zsGreen (1:1000, Clontech), STEM121 (1:500, Takara), TUJ1 (1:500, Biolegend), MAP2 (1:500, Sigma Aldrich), DARPP32 (1:500, Invitrogen), GAD65/67 (1:500, Abcam), GABA (1:500, Invitrogen), GFAP (1:500, Sigma) and Ki67 (1:500, Abcam). The species-appropriate Alexa Fluor™ secondary antibodies (1:500; Invitrogen) were used for visualisation of the primary antibody. DAPI (1:1000, Thermo Fisher Scientific) was used to confirm individual cell nuclei. Imaging was undertaken on a Nikon TE2000E inverted microscope (Nikon) equipped with a Nikon DS-Ri2 camera or on a Zeiss LSM 710 inverted confocal scanning laser microscope (Biomedical imaging Resource Unit, University of Auckland). Quantification of the number of TUJ1+, MAP2+, DARPP32+, GAD65/67+ or GABA+ hiLGEP-derived neurons was undertaken manually in ImageJ as a proportion of zsGreen+ or STEM121+ cells from a minimum of 200 zsGreen+ or STEM121+ cells.

### Statistical analysis

Statistical analyses were performed using IBM SPSS Statistics v28 (IBM Corporation). Levene’s test for equality of variances was performed on all data. A one-way or two-way analysis of variance was used for comparison of media composition and/or cell line. Post-hoc analyses were performed with the Bonferroni test. A two-way mixed ANOVA with subsequent simple main effect analysis using Bonferroni correction was used to compare the spontaneous exploratory forelimb use of hiLGEP-transplanted animals to sham overtime. All data are presented as mean ± SEM. Results were considered significant if p < 0.05. Data was plotted in GraphPad Prism v9.02.

## Results

### Adult human dermal fibroblasts can be directly reprogrammed to a lateral ganglionic eminence phenotype

Previous studies have demonstrated the requirement to drive hESCs to a LGEP fate prior to transplantation to promote *in vivo* differentiation to MSNs [15–19]. Following these findings, we refined our *SOX2/PAX6* cmRNA direct-to-hiNPC reprogramming protocol [22] to promote the generation of neural precursor cells with a hiLGEP phenotype and enhance differentiation to MSNs (Figure 1).

We first investigated the effect of adding the broad-spectrum protein kinase C inhibitor Gö6983 (5µM), the p160ROCK inhibitor Y-27632 (10µM) and 1% N-2 supplement (combination abbreviated to GYN), and 10µM retinoic acid to our standard NBA-based reprogramming medium containing 1mM valproic acid, 1% penicillin-streptomycin-glutamine, 2% B-27 without retinoic acid, 20ng/ml EGF, 20ng/ml FGF2, and 2µg/ml heparin. Most importantly, we included Activin A (25 ng/ml) to the media after 7 days of reprogramming. Activin A has been shown by Arber and colleagues [19] to induce a LGEP fate from hESCs and hiPSCs in a SHH-independent manner. After 14 days of reprogramming, we observed that NBA-based reprogramming medium supplemented with Activin A either with or without GYN had little effect on the expression of the striatal transcription factor *DLX2* compared to NBA-based reprogramming medium alone (Figure 2A). However, NBA-based reprogramming medium supplemented with Activin A up-regulated the expression of the lateral ganglionic eminence (LGE)-selective gene *CTIP2* (3.44 ± 1.46 fold), which was further up-regulated with the addition of GYN (11.96 ± 6.6 fold) (Figure 2A).

**Figure 2:**
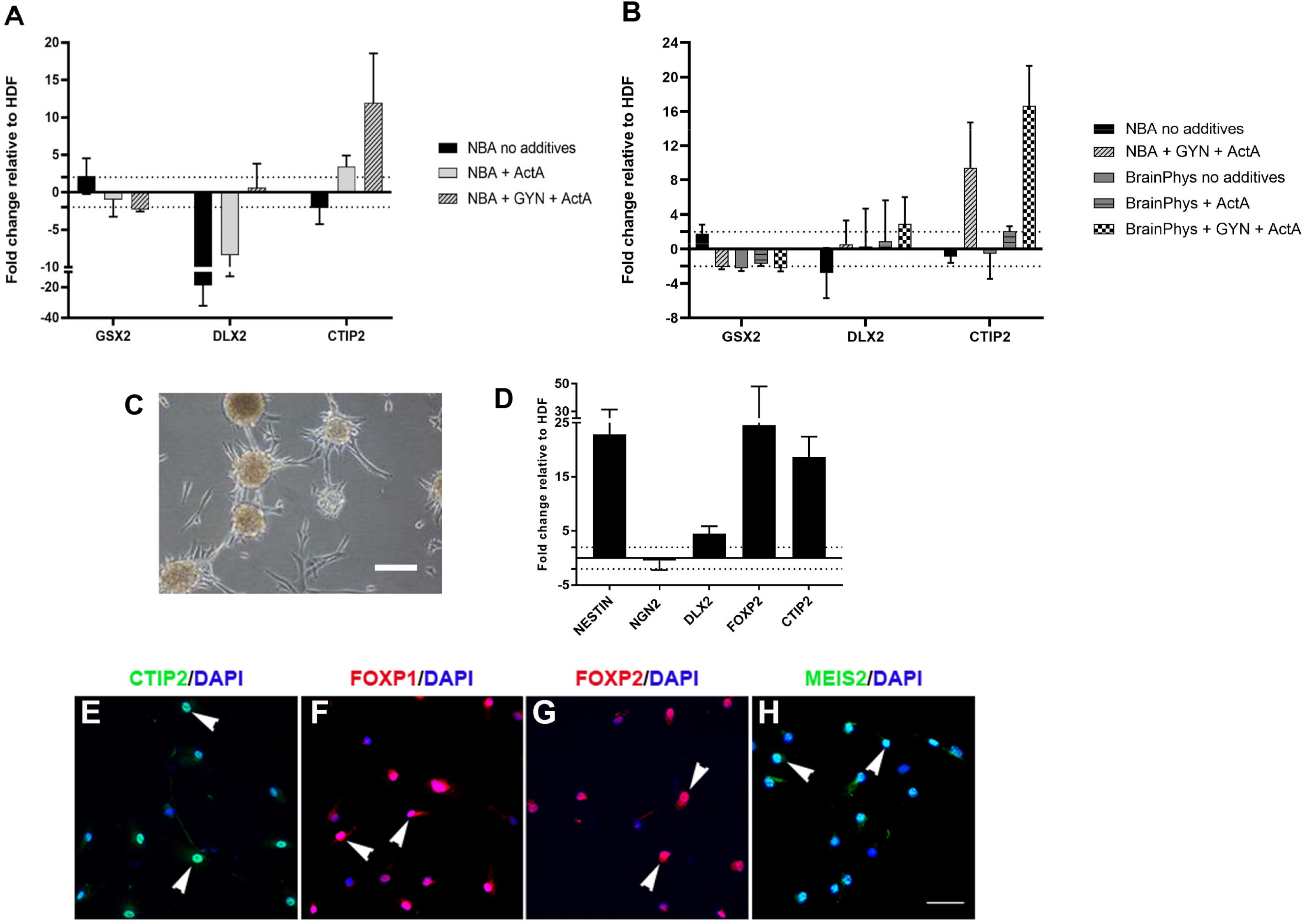
The effect of Activin A (ActA) with or without Gö6983, Y27632 and N-2 (GYN), and the effect of media on lateral ganglionic eminence (LGE) identity following 14 days of reprogramming. Graphs demonstrating the effect of NBA (A) or BrainPhys™ (B) media and ActA with or without GYN on *DLX2* and *CTIP2* expression. Data represent fold changes in mRNA expression relative to HDFs with mean ± SEM and n = 3 independent cell lines (1838, 2116 & 2298). Statistical significance was determined by one-way ANOVA with Bonferroni post-hoc test, * for p < 0.05. (C) Image of hiLGEP cultures after 14 days of reprogramming in BrainPhys™ supplemented with ActA and GYN. (D) Graph demonstrating expression of select transcription factors to confirm LGE identity after 14 days reprogramming in BrainPhys™ supplemented with ActA and GYN. Data represent fold changes in mRNA expression relative to HDFs with mean ± SEM and n = 6 independent cell lines. (E – I) Images of striatal marker expression to confirm LGE identity after 14 days reprogramming in BrainPhys™ supplemented with ActA and GYN. Scale bar: C = 100 µm; E – H = 50 µm.

We next compared the effect of the base medium BrainPhys™ to NBA on hiLGEP gene expression after 14 days reprogramming (Figure 2B). As BrainPhys™ is more representative of the central nervous system extracellular environment, we proposed it would enhance LGEP induction compared to NBA. After 14 days of reprogramming, we did not see an up-regulation of *CTIP2* in cells reprogrammed in BrainPhys™-based reprogramming medium without Activin A nor when supplemented with Activin A alone. However, when BrainPhys™-based reprogramming medium was supplemented with both GYN and Activin A we observed an up-regulation of *CTIP2* (16.67 ± 4.65 fold; Figure 2B). Statistical analysis revealed that *CTIP2* expression was significantly up-regulated when GYN and Activin A were both included in BrainPhys™-based reprogramming medium, compared to all other media (one-way ANOVA with Bonferroni post-hoc test, p = 0.005 compared to NBA, p = 0.033 compared to NBA + GYN + ActA, p = 0.011 compared to BrainPhys, and p = 0.015 compared to BrainPhys + ActA). Combined, these results indicate that Activin A in combination with Gö6983, Y-27632 and N-2 promotes the expression of the key LGEP gene *CTIP2*.

To confirm the ability of BrainPhys™ supplemented with GYN and Activin A to induce the expression of LGE transcription factors we collected cells after 14 days of reprogramming with *SOX2/PAX6* cmRNA and assessed both gene and protein expression. After 14 days of reprogramming, we observed formation of a network of progenitor (Figure 2C). At the gene level, there was an up-regulation of the pro-neural factor *NESTIN* and the LGE transcription factors *DLX2, FOXP2* and *CTIP2* when compared to aHDF (Figure 2D). We did not see a change in expression of the pro-glutamatergic neural factor *NGN2* [30]. At the protein level, the reprogrammed cells expressed 74.71% ± 3.12% CTIP2, 60.05% ± 6.38% FOXP1, 77.17% ± 1.19% FOXP2 and 85.01% ± 1.03% MEIS2 (Figures 2E-I). Based on these findings we propose BrainPhys™-based reprogramming medium supplemented with GYN and Activin A promotes the induction of an LGE precursor fate.

### Directly reprogrammed human lateral ganglionic eminence precursors differentiate to functional DARPP32-positive neurons in vitro

To ensure optimal differentiation of hiLGEPs to an MSN fate, we compared the effect of NBA- and BrainPhys™-based media on neuronal differentiation. We also investigated whether the addition of Activin A for the first 7 days of differentiation could enhance striatal differentiation (Figure 1). hiLGEPs that were reprogrammed in NBA-based medium were subsequently differentiated in NBA-based medium and hiLGEPs reprogrammed in BrainPhys™-based medium were differentiated in BrainPhys™-based medium. We observed that differentiation of hiLGEPs in NBA-based striatal differentiation medium containing Activin A resulted in 1.2% ± 0.33% to 14.33% ± 3.4% TUJ1/DAPI-positive cells regardless of the reprogramming protocol (Figure 3A). In contrast, hiLGEPs reprogrammed in BrainPhys™ supplemented with GYN and Activin A and differentiated in BrainPhys™-based striatal differentiation medium containing Activin A significantly enhanced the generation of TUJ1-positive cells to 63.8% ± 4.59% of the total DAPI+ cell population compared to 9.4% ± 1.74% TUJ1/DAPI-positive cells from hiLGEPS reprogrammed with BrainPhys™ and Activin A alone (one way ANOVA with Bonferroni post-hoc test, p = 1.83E-16 – 9.7E-14 for BainPhys +GYN +ActA compared to all other conditions; Figure 3A). Similarly, the number of DARPP32-positive cells following differentiation in NBA-based striatal differentiation medium with Activin A was very low (0.15% ± 1.15% to 10.76% ± 2.2%; Figure 3B). However, hiLGEPs reprogrammed in BrainPhys™ supplemented with GYN and Activin A and differentiated in BrainPhys™ -based striatal differentiation medium containing Activin A resulted in 42.45% ± 2.72% of the total DAPI+ cell population expressing DARPP32 compared to 3.77% ± 0.95% of DARPP32/DAPI-positive cells from hiLGEPS reprogrammed with BrainPhys™ and Activin A alone (one-way ANOVA with Bonferroni post-hoc test, p = 0.001 for NBA +ActA, p = 0.004 for NBA +GYN +ActA, p = 0.0498 for BrainPhys +ActA and p = 1.41E-14 for BrainPhys +GYN +ActA all compared to NBA no additives; Figure 3B). The findings demonstrate that reprogramming *SOX2/PAX6* cmRNA-transfected aHDFs in BrainPhys™ supplemented with GYN and Activin A promotes the induction of hiLGEPs and differentiation of hiLGEPs in BrainPhys™ media supplemented with Activin A enhances the generation of TUJ1-positive and DARPP32-positive neurons. To confirm the robustness of these findings, we investigated the expression of the mature human neuron protein PGP9.5 in hiLGEP-derived neurons by MACS MicroBead magnetic cell sorting (Figure 3C). hiLGEPs reprogrammed in NBA-based medium with GYN and Activin A gave rise to 9.89% ± 0.088% neurons of the total cell population expressing PGP9.5, while hiLGEPs reprogrammed in BrainPhys™ medium supplemented with Activin A alone gave rise to 18.8% ± 3.34% neurons of the total cell population expressing PGP9.5. In contrast, hiLGEPs reprogrammed in BrainPhys™ medium with GYN and Activin A yield the largest proportion of neurons expressing PGP9.5 from the total differentiated cell population with 63.04% ± 3.2% PGP9.5 expression (one-way ANOVA with Bonferroni post-hoc test, p = 0.307 for NBA +GYN +ActA compared to BrainPhys +ActA, p = 0.000078 for NBA +GYN +ActA compared to BrainPhys +GYN +ActA and p = 0.000223 for BrainPhys +ActA compared to BrainPhys +GYN +ActA; Figure 3C). Taken together, these results support our claim that *SOX2/PAX6* cmRNA-transfected aHDFs reprogrammed in BrainPhys™ medium supplemented with GYN and Activin A promotes the acquisition of a hiLGEP fate which further differentiate into high yields of TUJ1-positive, PGP9.5-positive and DARPP32-positive neurons in BrainPhys™ media supplemented with Activin A.

**Figure 3:**
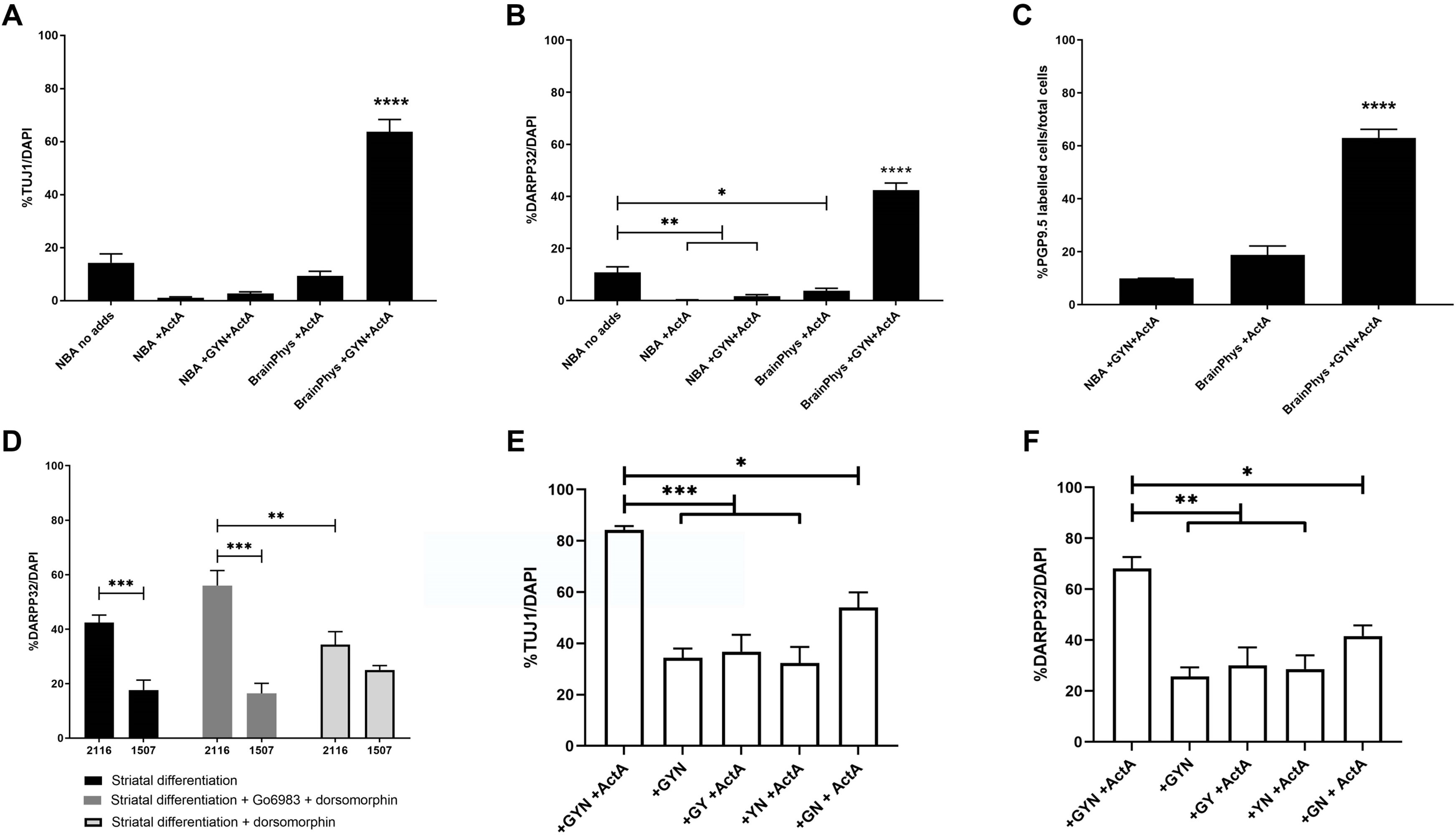
The effect of media and dorsomorphin with or without Gö6983 on the differentiation of hiLGEPs to striatal neurons. Graphs demonstrating the effect of NBA or BrainPhys™ media on the proportion of TUJ1- (A) or DARPP32- (B) positive cells after 14 days differentiation. Data represent mean ± SEM with n = 3 independent cell lines (1507, 2116 and 2298; 8 fields of view and a minimum of 500 DAPI+ cells per line). Statistical significance determined by one-way ANOVA with Bonferroni post-hoc test, * for p < 0.05; ** for p < 0.01; **** for p < 0.0001. (C) Graph demonstrating the effect of NBA or BrainPhys™ reprogramming media with Activin A, and with or without GYN on the proportion of PGP9.5-labelled cells after 14 days of differentiation. Data represent mean ± SEM with n = 3 independent cell lines (1507, 2116 and 2298; 12 replicate wells per line pooled for MACS® MicroBead magnetic cell sorting). (D) Graph demonstrating the effect of dorsomorphin with or without Gö6983 on the proportion of DARPP32-positive cells compared between 2 independent cell lines (2116 & 1507; n = 8 replicate wells from independent experiments /line). Statistical significance determined by two-way ANOVA with simple main effects, ** for p < 0.01, *** for p < 0.001. Graphs demonstrating the effect of reprogramming with GYN alone or in combination with ActA with or without all three GYN compounds on the proportion of (E) TUJ1-positive cells and (F) DARPP32-positive cells after 14 days of differentiation. Data represent mean ± SEM with n = 3 replicate wells from independent experiments but the same donor cell line (2116). Statistical significance determined by one-way ANOVA with Bonferroni post-hoc test, * p ≤ 0.05; ** p ≤ 0.01; *** p ≤ 0.001.

Finally, we investigated whether supplementation of BrainPhys™-based striatal differentiation medium with the AMP-activate kinase inhibitor dorsomorphin for the first 5 days of differentiation or supplementation with the broad-spectrum protein kinase C inhibitor Gö6983 for the first 5 days of differentiation followed by dorsomorphin for the next 5 days of differentiation could further enhance the generation of DARPP32-positive neurons. We also compared the effect of dorsomorphin with or without Gö6983 between 2 independent cell lines (2116 and 1507) to assess consistency and robustness of DARPP32-positive neuronal yield. Interestingly the addition of dorsomorphin with or without Gö6983 did not significantly alter the proportion of DARPP32-positive neurons compared to standard BrainPhys™-based striatal differentiation medium (Figure 3D). There was a significant interaction of differentiation condition and cell line by two-way ANOVA (F(2,42) = 7.696, p = 0.001) indicating the different media conditions had an effect on DARPP32 yield which differed between cell lines. While a significant increase in DARPP32-positive cells was seen when striatal differentiation medium was supplemented with Gö6983 and dorsomorphin compared to dorsomorphin alone for line 2116 (p = 0.007; Figure 3D), further post hoc analysis determined that BrainPhys™ - based striatal differentiation medium supplemented with dorsomorphin alone produced the most consistent DARPP32-positive cell yield between cell lines (p = 0.088 for 2116 compared to 1507 in striatal differentiation + dorsomorphin; p = 0.0008 for 2116 compared to 1507 in striatal differentiation only; p = 0.0003 for 2116 compared to 1507 in striatal differentiation + Gö6983 + dorsomorphin). Based on these findings, we chose to use BrainPhys™ -based striatal differentiation medium supplemented with dorsomorphin for the first 5 days and Activin A for the first 7 days of differentiation.

To assess the contribution and need for each of the reprogramming compounds to generate high yields of DARPP32/TUJ1-positive hiLGEP-derived neurons, we considered the neuronal yield of hiLGEPs reprogrammed with GYN alone or in combination with Activin A, with or without all three GYN compounds (Figure 3E & F). We found that when GYN and Activin A were added in combination to the reprogramming medium the hiLGEPs gave rise to 84.22% ± 1.25% TUJ1-postive cells which was a significant increase in yield when compared to all other media conditions (p = 0.013 when compared to +GN +ActA and p = 0.000236 to 0.00491 when compared to all other conditions; Figure 3E). Similarly, when GYN and Activin A were added in combination to the reprogramming medium the hiLGEPs gave rise to 68.20% ± 3.64% DARPP32-postive cells which was a significant increase in yield when compared to all other media conditions (p = 0.026 when compared to +GN +ActA and p = 0.001 to 0.002 when compared to all other conditions; Figure 3F).

Using these culture conditions, we were able to generate 66.59% ± 1.66% TUJ1+/DAPI+ neurons, 53.18% ± 3.21% DARPP32+/DAPI+ neurons, 56.26% ± 1.65% GABA+/DAPI+ neurons and 48.12% ± 6.12% GAD65/67+/DAPI+ neurons (n = 4 independent cell lines), which exhibited extensive neurite outgrowth and network formation (Figure 4A – E). To confirm the generation of a medium spiny striatal neuron fate, we examined the co-expression of these markers and found 70.18% ± 7.14% DARPP32+/TUJ1+ neurons, 50.69% ± 5.09% GABA+/TUJ1+ neurons, 39.73% ± 4.09% DARPP32+/GABA+ neurons and 38.37% ± 5.03% DARPP32+/ GAD65/67+ neurons (Figure 4F – H). Functionality of hiLGEP-derived neurons was demonstrated by live-cell calcium imaging (Supplementary Figure 2). Cultures were loaded with the fluorescence-based calcium indicator Cal-520, and exposed to either 0µM, 12.5µM, 25µM, 37.5µM, or 50µM glutamate. Cal-520 fluorescence was measured as the percentage increase in average fluorescence intensity relative to baseline at time 0 and normalised to the average fluorescence intensity in response to no glutamate. hiLGEP-derived neurons exhibited an increase in Cal-520 fluorescence with increasing concentration of glutamate, with the average intensity greatest for 25µM glutamate peaking at 90 seconds after glutamate administration followed by a reduction in average intensity between 95 - 100 seconds of -300 to -750 % relative to baseline (Figure 4I & J). hiLGEP-derived MAP2+ neurons were also shown to co-express the pre-synaptic marker SYN1 and the post-synaptic marker PSD-95 (Figure 4K & L), indicating synapse formation crucial to the functionality of the neuronal network.

**Figure 4:**
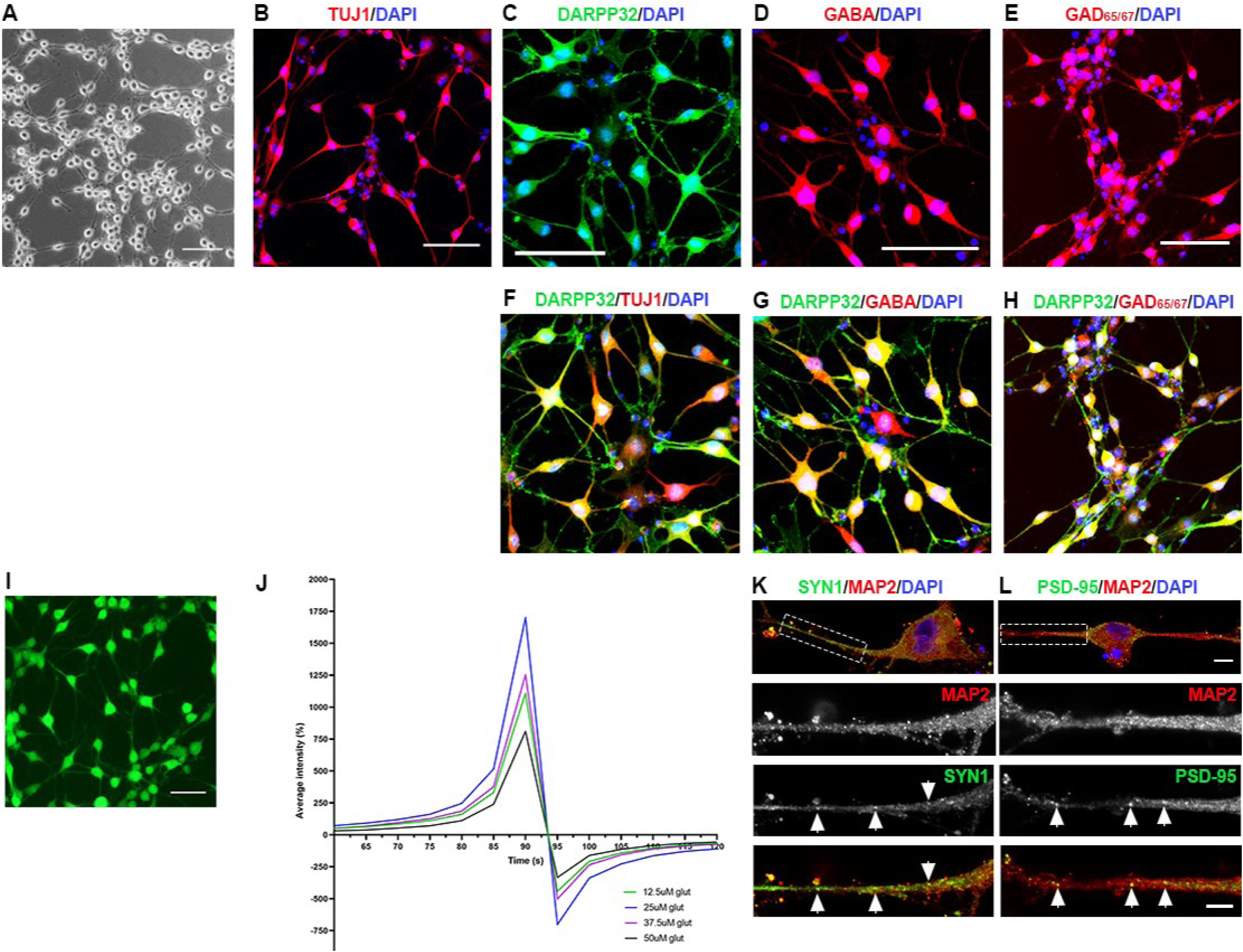
Differentiation of hiLGEPs to striatal neurons. (A) Bright-field image of hiLGEP-derived neurons after 14 days of differentiation in BrainPhys™ media supplemented with dorsomorphin. Images of (B) TUJ1-positive neurons, (C) DARPP32-positive neurons, (D) GABA-positive neurons, (E) GAD65/67-positive neurons, and DARPP32-positive neurons co-expressing (F) TUJ1, (G) GABA and (H) GAD65/67 after 14 days of differentiation in BrainPhys™ media supplemented with dorsomorphin. (I) Image of Cal-520 stain in hiLGEP-derived neurons after 14 days of differentiation in BrainPhys™ media supplemented with Activin A and dorsomorphin. Scale bars: 100 µm. (J) Cal-520 fluorescence was measured as the percentage increase in average fluorescence intensity relative to baseline fluorescence at time 0 and normalised to the average fluorescence intensity measured in response to no glutamate. Percentage increase in average fluorescence intensity was measured for 120 seconds in the presence of no glutamate, 12.5µM, 25µM, 37.5µM and 50µM glutamate; n = 75 cells. Images of MAP2-positive neurons co-expressing (K) the pre-synaptic marker SYN1 and (L) the post-synaptic marker PSD-95 after 14 days of differentiation in BrainPhys™ media supplemented with Activin A and dorsomorphin. Scale bar: 10µm and 5µm for higher magnification images. Arrows indicate SYN1+ and PSD-95+ puncta.

### Transplantation of directly reprogrammed human lateral ganglionic eminence precursors reduces impairment of spontaneous exploratory forelimb use

The spontaneous exploratory forelimb use test is a non-drug induced test of forelimb locomotor function that is dependent on the integrity of intrinsic striatal neurons, the nigrostriatal dopaminergic system and the sensorimotor area of the neocortex. Following a unilateral striatal lesion, rats will preferentially use the forelimb ipsilateral to the lesion to initiate and terminate weight-shifting movements during rearing and exploration along vertical surfaces [26, 27]. Using the spontaneous exploratory forelimb use test, we investigated the effect of transplanting hiLGEPs into the QA-lesioned striatum on forelimb locomotor function at 2-, 4-, 12- and 14-weeks following transplantation (Figure 5A).

**Figure 5:**
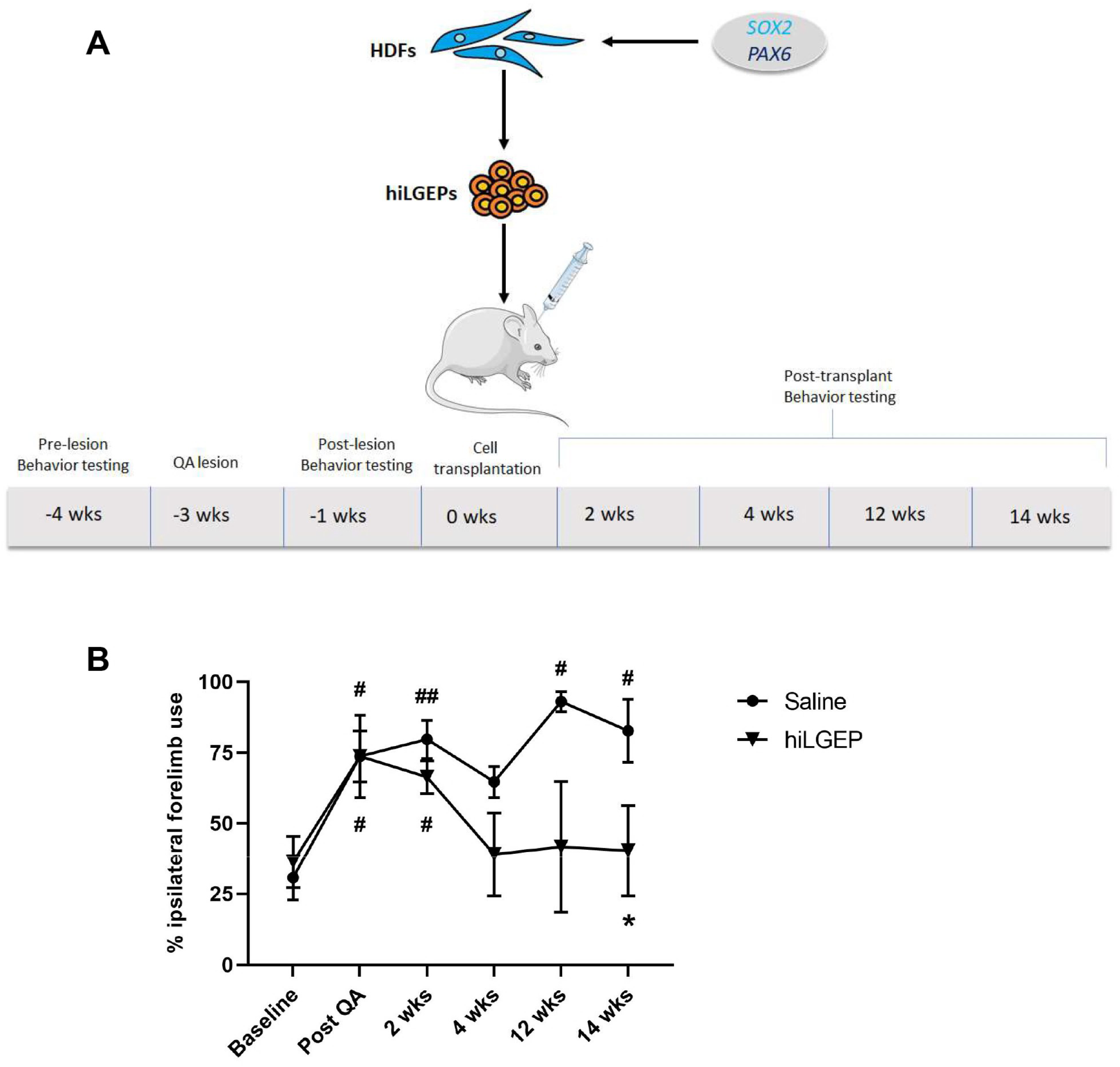
Transplantation of hiLGEPs significantly reduces impairment of forelimb locomotor function. (A) Schematic diagram demonstrating the experimental time-line of QA lesioning, hiLGEP transplantation and behaviour testing. (B) Rats were tested at weeks 2, 4, 12 and 14 following transplantation for spontaneous exploratory forelimb use. Saline-treated rats exhibited a significant increase in preference for the use of the ipsilateral forelimb over time compared to baseline (# for p < 0.05, ## for p < 0.01). In contrast, rats transplanted with hiLGEPs only exhibited a significant increase in preference for the use of the ipsilateral forelimb following QA lesioning and 2 weeks post-transplantation compared to baseline (# for p < 0.05). At 14 weeks post-transplantation, a significant difference in the preferential use of the ipsilateral forelimb was observed in rats transplanted with hiLGEPs compared to post-QA (* for p < 0.05). Data represents mean ± SEM. Statistical significance was determined by a two-way mixed ANOVA with subsequent simple main effects analysis. This figure was partly generated using Servier Medical Art, provided by Servier, licensed under a Creative Commons Attribution 3.0 unported licence.

A two-way mixed ANOVA with subsequent simple main effects analysis demonstrated a significant interaction between time and treatment (F = 2.83, df = 5, p = 0.043). After QA lesioning, both treatment groups exhibited a significant preference for use of the ipsilateral forelimb compared to baseline (saline treatment, p = 0.017; hiLGEP treatment, p = 0.027, Figure 5B). Animals that received saline treatment continued to exhibit a significant preference for ipsilateral forelimb use over time when compared to baseline (2 weeks post-transplant, p = 0.008; 12 weeks post-transplant, p = 0.036; 14 weeks post-transplant, p = 0.026; Figure 5B). In contrast, hiLGEP transplanted animals only exhibited a significant preference for ipsilateral forelimb use at 2 weeks post-transplant when compared to baseline (p = 0.04; Figure 5B). Extending this observation, a significant decrease in the ipsilateral forelimb use of hiLGEP transplanted animals was seen at 14 weeks post-transplant when compared to after QA lesioning (p = 0.027; Figure 5B). These results indicate that striatal transplantation of hiLGEPs can reduce impairment of spontaneous exploratory forelimb use caused by QA lesioning of the striatum.

### Human induced lateral ganglionic eminence precursors survive transplantation into the QA lesioned striatum and differentiate to medium spiny striatal neurons

We investigated the capability for *SOX2/PAX6* cmRNA directly reprogrammed hiLGEPs to survive transplantation and differentiate to medium spiny striatal neurons in the rat QA lesion model of HD. hiLGEP-derived neurons were identified in the QA lesioned rat striatum by expression of either zsGreen (8 weeks post-transplantation; Figure 6) or the human cytoplasmic marker STEM121 (14 weeks post-transplantation; Figure 7). At both 8 and 14 weeks after transplantation, zsGreen- or STEM121-positive cells were detected within a defined boundary in the anterior aspect of the striatum of hiLGEP transplanted animals (Figure 6A and 7A). zsGreen- and STEM121-positive cells displayed a distinctive neuronal morphology with extensive neurite outgrowth seen by 14 weeks post-transplant (Figure 7A&A’). We did not observe zsGreen- nor STEM121-positive cells exhibiting an astrocytic morphology and there was no co-expression of zsGreen or STEM121 with GFAP (Figure 6E). The transplanted hiLGEPs did not proliferate post-transplantation as seen through a lack of expression of the proliferation marker Ki67 in the zsGreen+ hiLGEPs (Figure 6F). At 14 weeks post-transplant we observed ∼ 23% of transplanted STEM121+ hiLGEPs remained in the QA-lesioned striatum.

**Figure 6:**
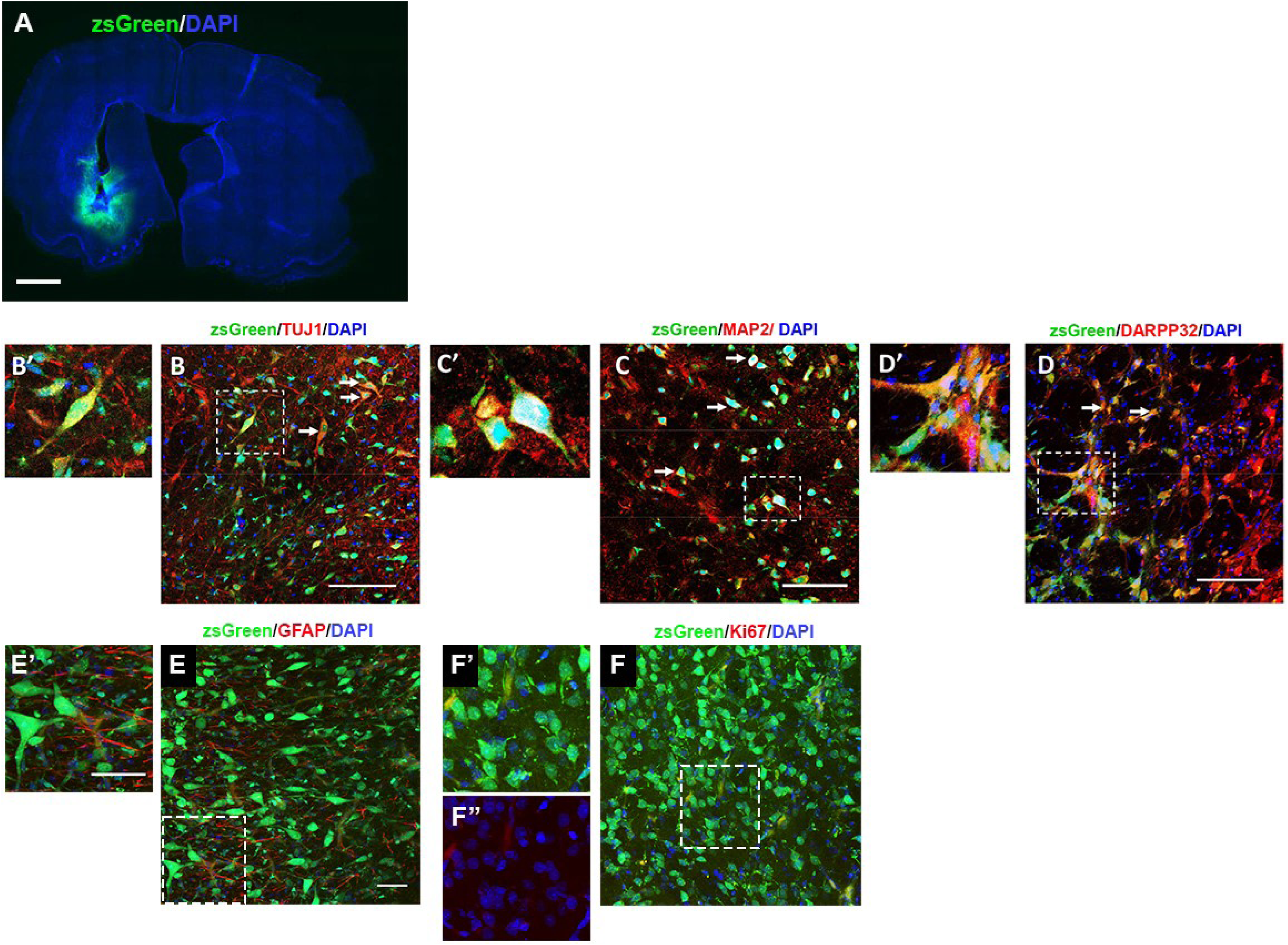
Transplanted hiLGEPs survive and generate DARPP32-positive cells 8 weeks following transplantation into the QA lesioned rat brain. (A) View of a whole section demonstrating the DAPI+ unlesioned and lesioned striatum and the zsGreen+ transplant core. Scale bar: 1mm. Images of zsGreen-labelled cells in the QA lesioned striatum co-expressing (B & B’) TUJ1, (C & C’) MAP2 and (D & D’) DARPP32 (arrows). B’,C’, D’, E’ and F’are high magnification images of cells indicated in B, C, D, E and F by boxed area. F’’ is a high magnification image representing Ki67 and DAPI co-staining only. Scale bars = 50 µm.

**Figure 7:**
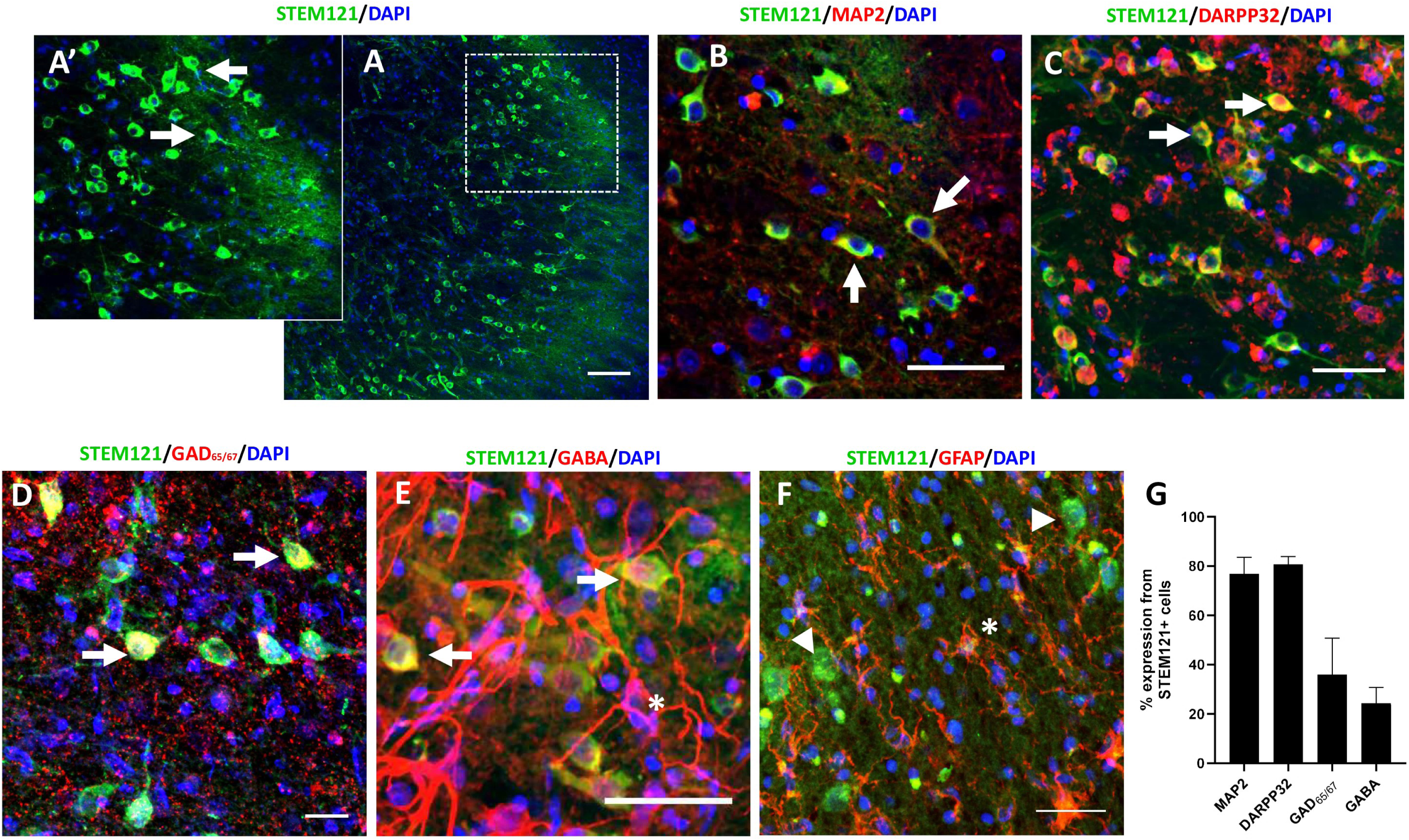
Transplanted hiLGEPs survive and generate medium spiny striatal neurons 14 weeks following transplantation into the QA lesioned rat brain. (A) Distribution of STEM121-positive cells in the QA lesioned striatum. (A’) High magnification image of A showing neuronal morphology of STEM121-positive cells with extensive neurite extensions (arrows). Confocal images demonstrating that STEM121-positive cells co-express the neuronal marker MAP2 (B) and the medium spiny striatal markers DARPP32 (C), GAD65/67 (D) and GABA (E); Examples indicated by arrows. (F) STEM121-positive cells do no co-express the astrocyte marker GFAP. Arrow heads indicate STEM121-positive/GFAP-negative cells. * indicates STEM121-negative GABA-positive or GFAP-positive astrocytic cell. Scale bars = 100µm (A), 50µm (B – E) and 30µm (F). (G) Graph summarising the % phenotypic marker from STEM121+ hiLGEP-derived neurons 14 weeks post-transplantation. Data represent mean ± SEM.

To determine the resultant phenotype of the hiLGEP-derived neurons 8 and 14 weeks after transplantation into the QA lesioned striatum, double-label immunofluorescence for zsGreen or STEM121 and cell type-specific markers was performed (Figures 6 & 7). Lack of zsGreen and STEM121 staining in saline animals and corresponding staining for MAP2, DARPP32, GABA, GAD65/67 and GFAP are provided in Supplementary Figure 3. We observed that 8 weeks after transplantation a population of hiLGEPs co-expressed zsGreen and the neuronal marker TUJ1 (Figure 6A & A’). Quantification of zsGreen-labelled cells showed that 37.8% ± 19.1% of zsGreen-labelled hiLGEP transplanted cells were positive for TUJ1.

Expression of the neuronal marker MAP2 was also observed 8 weeks after transplantation of hiLGEP cells with approximately 50.3% ± 9.4% of zsGreen-labelled-cells positive for MAP2 (Figure 6B&B’). To confirm the ability for transplanted hiLGEPs to generate MSNs, sections were co-labelled for zsGreen and DARPP32. In agreement with our *in vitro* findings, we observed a population of zsGreen-labelled cells expressing DARPP32 (66.3% ± 8.46%) 8 weeks post-transplantation into the QA lesioned striatum (Figure 6C&C’).

At 14 weeks post-transplantation, 76.8% ± 6.79% STEM121-positive cells co-expressed MAP2 (Figure 7B & G) while 80.7% ± 3.13% STEM121-positive cells co-expressed DARPP32 (Figure 7C & G). 35.9% ± 14.84% STEM121-positive cells co-expressed the enzyme GAD65/67 (Figure 7D & G) with 24.36% ± 6.34% STEM121-positive cells co-expressing GABA (Figure 7E & G). We also observed GABA-positive / STEM121-negative cells displaying an astrocytic morphology (Figure 7E). STEM121-positive cells did not co-express GFAP, rather GFAP-positive cells were exclusively STEM121-negative (Figure 7F). Astrocytes have been shown to synthesis and uptake GABA via multiple pathways [31, 32]. The lack of STEM121 co-expression indicates the astrocytes are from the host rat brain in response to the QA lesion. Furthermore, this observation combined with the lack of STEM121, and zsGreen, co-expression with GFAP demonstrates the transplanted hiLGEPs do not generate astrocytes following transplantation.

These results indicate that *SOX2/PAX6* cmRNA direct reprogramming generates hiLGEPs which survive transplantation and differentiate into medium spiny striatal neurons in the QA lesioned rat striatum.

## Discussion

This study demonstrates for the first time that aHDFs can be directly reprogrammed with *SOX2* and *PAX6* cmRNA to a hiLGEP phenotype which, following transplantation into the QA lesioned rat striatum, results in the generation of DARPP32- and GABA-positive neurons and restoration of motor function. One of the key aspects of this study is the use of cmRNA to generate hiLGEPs for transplantation. Regarding both safety and efficiency, mRNA provides an ideal non-viral, non-integrating delivery system for cell reprogramming [24, 25, 33]. The cmRNA system used in this study allows for mRNA transfection without immune response inhibition through the replacement of uridine and cytidine residues with chemically modified uridine and cytidine analogues, respectively, reducing the activation of an innate immune response and increasing mRNA stability [34]. As such, the use of cmRNA to generate reprogrammed donor cells for cell replacement therapy is highly attractive as it provides an efficient and stable system of gene delivery without the risk of genomic integration and insertional mutation inherent to all DNA-based methodologies as well as allowing cell reprogramming without residual tracers of transgenes [24]. We propose these features make cmRNA an excellent option for the clinical translation of reprogramming-based cell replacement therapies.

Another key feature of this study is the demonstration that aHDFs can be directly reprogrammed to a LGEP fate. Previous studies using either hESC or hiPSCs have demonstrated the requirement to transplant lineage-specific neural precursor cells to ensure complete differentiation to a striatal phenotype [15–19, 35]. Through knowledge in human development, previous protocols have employed a defined dose of SHH or SHH together with pharmacological inhibition of Wnt signalling, to obtain LGE-like neural precursors from hESCs and hiPSCs [16, 17, 35–37]. In contrast, Arber and colleagues [19] reported that Activin A induces LGE characteristics to hESC- and hiPSC-derived neural precursors which readily give rise to DARPP32-expressing neurons in culture and following transplantation into the QA lesion rat model of HD. Activin A is a multifunctional member of the TGFβ family that has been shown to induce forebrain neurogenesis in a neuronal subtype-specific manner [38, 39]. Activin A receptors and the effector protein Smad2 are expressed in the developing LGE and functionally interact with Dlx to induce the development of telecenphalic GABAergic neurons [40]. Interestingly, the pattern of gene expression induced by Activin A is different from that induced by SHH, with SHH eliciting a strong inductive effect on medial ganglionic eminence (MGE) regulator genes such as *NKX2.1* and genes expressed in both the MGE and LGE (*DLX2, GSX2)* in a dose-dependent manner. In contrast, the expression of LGE-specific genes (e.g. *CTIP2, FOXP1/2)* changes very little in response to SHH [19, 41–44]. Arber and colleagues [19] therefore proposed that the main role for SHH is to suppress dorsal fate rather than to ‘direct’ a striatal fate. In contrast, Activin A may act directly to bias the differentiation of neural precursors towards an LGE/striatal phenotype [19].

Therefore, adapting the protocol developed by Arber and colleagues [19] we have demonstrated that culture of aHDFs in Activin A combined with Gö6983, Y27632 and N-2 (GYN) following *SOX2/PAX6* cmRNA direct reprogramming results in neural precursor cells expressing the striatal factors *DLX2*, FOXP1, FOXP2, CTIP2 and MEIS2. Based on this profile, and in particular the significant up-regulation of *CTIP2* expression, our findings support the use of Activin A to induce a LGE fate which was further promoted by the addition of Gö6983, Y27632 and N-2. The generation of hiLGEPs by direct reprogramming was further confirmed by the generation of DARPP32-positive neurons following *in vitro* differentiation of hiLGEPs in BrainPhys™ -based striatal differentiation medium supplemented with Activin A and dorsomorphin. The hiLGEP-derived striatal neurons were shown to be functional by calcium imaging and displayed synapse formation through the expression of the pre- and post-synaptic markers SYN1 and PSD-95, respectively.

Following the establishment of a LGE fate we investigated the ability for hiLGEPs to survive transplantation, differentiate to MSNs and improve motor function in the QA lesion rat model of HD. Eight weeks following transplantation ∼66% of zsGreen-labelled hiLGEPs had differentiated to express the MSN marker DARPP32 within the QA lesioned striatum and by 14 weeks post-transplantation ∼81% of STEM121+ hiLGEPs co-expressed DARPP32. This study is the first to investigate the transplantation of directly reprogrammed hiLGEPs and as such there is no study to compare our results with. Furthermore, very few papers investigating the survival and differentiation of transplanted hESC- or hiPSC-derived NPCs in the QA lesioned striatum have provided quantification of DARPP32-positive neuronal generation [9]. Arber and colleagues [19] reported low numbers of DARPP32-positive cells 8 weeks following transplantation of hESC-derived LGEP-like precursor cells into the QA lesioned striatum and it wasn’t until 16 weeks post-transplantation that ∼50% DARPP32 positive neurons were detected *in vivo*. Furthermore, Besusso and colleagues [35] recently demonstrated that only 2-3% of hESC-derived striatal progenitors generated DARPP32-positive neurons 8 weeks post-transplant into the QA lesioned striatum with 45% of transplanted cells expressing CTIP2 only. While Besusso and colleagues [35] did not use Activin A to induce a LGE phenotype prior to transplant, the difference in DARPP32 yield may indicate that directly reprogrammed hiLGEPs are more responsive to differentiation cues within the QA lesioned striatum than hESC generated precursor cells.

The effect of the environment on neural differentiation is well known [45–47] with a dynamic interplay occurring between injury-mediated maturation signals and those maintaining an immature neural phenotype that are neurogenic and niche-derived [48–50]. Cell loss has been shown to promote neural differentiation and maturation through the expression of molecular controls [49–51]. Furthermore, the QA lesioned environment is known to support cell survival of primary striatal grafts to a greater extent than the unlesioned striatum, and may enhance lineage-specific differentiation and maturation [52, 53] through injury-induced expression of neurotrophic factors [54].

Tumour formation post-transplant is one concern that requires mitigating for cell transplantation strategies to progress to the clinic. This direct reprogramming to hiLGEP protocol bypasses the formation of pluripotent intermediates and utilises stable, non-immunogenic and non-integrating cmRNA, a clinically viable reprogramming method. What is more, hiLGEPs did not proliferate post-transplant *in vivo* as seen by a lack of Ki67 expression.

Most importantly, this study demonstrates that transplantation of directly reprogrammed hiLGEPs to the QA lesioned striatum can restore motor function impairment as determined by spontaneous exploratory forelimb use when compared to saline treated animals. Again, very few studies have investigated the behavioural effect of hESC- or hiPSC-derived NPCs following transplantation into either the QA lesion rat model or transgenic mouse models of HD [9]. In the QA lesion model, Carri and colleagues [15] observed a reduction in apomorphine-induced rotations 3 weeks following transplantation of hESC-derived NPCs while Arber and colleagues [19] reported no change in apomorphine rotational asymmetry following hESC-derived LGEP transplantation. Besusso and colleagues [35] however demonstrated a significant improvement in both the Vibrissae-evoked Hand Placing test and the Adjusting Steps test 4 weeks following transplantation of hESC-derived NPCs into the QA lesioned striatum. In addition, Jeon and colleagues [20] reported significant improvement in the Stepping and Staircase tests from 4 weeks post-transplantation as well as a reduction in apomorphine-induced rotations 6 weeks following transplantation of hiPSC-derived NPCs in the QA lesion model. In contrast to previous studies, we did not observe an improvement in spontaneous exploratory forelimb use until 14 weeks following transplantation of hiLGEPs into the QA lesioned striatum. However, while not significant compared to saline-treated animals, the percentage of ipsilateral forelimb use for hiLGEP transplanted animals had returned to baseline levels by 4 weeks post-transplantation, consistent with the motor function improvement seen with hESC- and hiPSC-derived NPC transplants.

An ongoing issue that requires addressing for cell transplantation therapy to be a viable long-term therapy for HD, is the propagation of mutant huntingtin (mHTT) from the host into the grafted cells. Indeed, HD has been described as a prion disease, meaning that mHTT aggregates can propagate from cell to cell through functional networks, resulting in non-cell autonomous damage [55–57]. It is warranted for future cell transplantation studies to include investigation of mHTT propagation into the grafted cells and the therapeutic benefit of blocking this propagation to predict the long-term therapeutic benefit of the given cell transplantation strategy.

In conclusion, this study provides proof of concept and demonstrated that aHDFs can be directly reprogrammed to hiLGEPs following transient cmRNA-mediated over-expression of *SOX2* and *PAX6* and exposure to Activin A, Gö6983, Y-27632 and N-2 in a BrainPhys™-based reprogramming medium. We have demonstrated for the first time that transplantation of directly reprogrammed hiLGEPs results in a high yield of medium spiny striatal neurons in the QA lesioned striatum in agreement with the observations made by Arber and colleagues [19]. Furthermore, transplantation of hiLGEPs in the QA lesioned striatum significantly reduces motor function impairment as determined by spontaneous exploratory forelimb use when compared to saline treated animals. Based on these findings, we propose the use of cmRNA directly reprogrammed hiLGEPs offers an effective and clinically viable strategy for cell replacement therapy to treat HD.

## Declarations

### Ethics Approval

All animal procedures strictly complied with the University of Auckland Animal Ethics Guidelines, in accordance with the New Zealand Animal Welfare Act 1999 and international ethical guidelines. All efforts were made to minimize the number of animals used and their suffering. AEC R21614, approved 03/05/2021, for project entitled “Novel Cell Replacement Therapy for Huntington’s Disease”.

### Consent for publication

All authors have provided written consent for publication.

### Availability of data and materials

The datasets used and/or analysed during the current study are available from the corresponding author on reasonable request.

### Competing interests

The authors declare no competing interests except for a provisionally filed patent International PCT filing AU2023902064; Cellular Reprogramming. Connor, B and Samuel, A.J.

### Funding

This research was supported by the Health Research Council of New Zealand, Auckland UniServices Limited Inventor Fund, the Maurice and Phyllis Paykel Trust and the Hudson-Nilon Medical Research Trust.

### Author Contributions

Amy McCaughey-Chapman; Conception and design, Collection and/or assembly of data, Data analysis and interpretation, Manuscript writing, Final approval of manuscript.

Anne Lieke Burgers; Collection and/or assembly of data, Data analysis and interpretation, Final approval of manuscript.

Catharina Combrinck; Collection and/or assembly of data, Data analysis and interpretation, Final approval of manuscript.

Laura Marriott; Collection and/or assembly of data, Final approval of manuscript. David Gordon; Collection and/or assembly of data, Final approval of manuscript.

Bronwen Connor; Conception and design, Financial support, Data analysis and interpretation, Manuscript writing, Final approval of manuscript

## Acknowledgements

Dr Carsten Rudolph and Dr Johannes Geiger, Ethris GmbH for provision of the *SOX2* and *PAX6* chemically modified mRNA.

---

**Supplementary Figure 1:** SOX2 and PAX6 expression reduces over the course of reprogramming and is no longer present in differentiated hiLGEPs. (A) Graph demonstrating the gene expression of *SOX2* and *PAX6* at Day 7 and Day 14 of reprogramming. Data represent fold changes in mRNA expression relative to HDFs with mean ± SEM and n = 3 independent cell lines. SOX2 and PAX6 protein expression in (B) reprogrammed hiLGEPs and (C) at Day 14 of differentiation. Scale bars: 100µm.

**Supplementary Figure 2:** Video demonstrating live-cell calcium imaging of hiLGEP-derived neurons loaded with the fluorescence-based calcium indicator Cal-520 and exposed 25µM glutamate.

**Supplementary Figure 3:** Immunohistochemical staining of saline-treated animals. Rats that received a saline injection show no expression of (A) zsGreen nor (B) STEM121, minimal expression of (C) MAP2, (D) DARPP32, (E) GAD65/67 and (F) GABA, and expression of (G) GFAP. Scale bars: 30µm,

## Notes

### Competing Interest Statement

The authors have declared no competing interest.

